# Sensitivity to Affective Touch Depends on Adult Attachment Style

**DOI:** 10.1101/373415

**Authors:** Charlotte Krahé, Mariana von Mohr, Antje Gentsch, Lisette Guy, Chiara Vari, Tobias Nolte, Aikaterini Fotopoulou

**Author notes:** Corresponding author: Mariana Von Mohr, University College London, CEHP Research Department 1-19 Torrington Place, London WC1E 7HB. Joint 1^st^ authorship.

## Abstract

Affective touch supports affiliative bonds and social cognition. However, it remains unknown whether pre-existing models of social relating influence the perception of affective touch. Here, we present the first study (*N*=44) to examine how individual differences in attachment styles relate to the perception of affective touch, as well as to a different non-social modality of interoception, namely cardiac perceived accuracy. Using the gold standard assessment of adult attachment (Adult Attachment Interview), we found that insecure attachment was associated with reduced pleasantness discrimination between affective vs. non-affective, neutral touch. Acknowledging the different traditions in measuring attachment, we also used a well-validated self-report questionnaire that pertains to explicit representations of current close relationships. Using this measure, we found that higher scores on an attachment anxiety dimension (but not an attachment avoidance) were associated with reduced pleasantness discrimination between affective vs. non-affective, neutral touch. Attachment patterns (in both measures) were not related to cardiac perception accuracy. These results corroborate and extend previous literature on the affectivity of touch and its relation with affiliative bonds and social cognition. Given that attachment was not related to perceived cardiac accuracy, these findings point to the specificity of the relationship between affective touch and attachment.

## Sensitivity to Affective Touch Depends on Adult Attachment Style

Attachment theory is one of the most influential theories of the development of close social relationships^1,2^. Its key tenet is that infants have an innate drive to form a close bond with their primary caregivers to ensure their survival and well-being in times of threat. In the past decades, the emphasis in attachment research has been influenced by the additional, cognitive hypothesis that differences in the responsiveness and availability of caregivers to the infant’s attachment needs lead to the development of internal working models (IWM) of social relating and associated affect regulation strategies^3^. These working models are described as affective-cognitive schemas, termed ‘attachment representations’ or most generally referred to as attachment styles, that are transferred from parental figures to romantic relationships^4^ and remain relatively stable across the life span^5^. For example, secure attachment is characterized by positive views of self and other, and the belief that one can turn to others for support and those others will be responsive^6^.

The emphasis on these IWM in attachment theory has somewhat shifted attention away from Bowlby’s original attention to physical ‘proximity seeking’ as the primary behavioural strategy for coping with threat (in a wider sense)^1,2^. Crucially, a central aspect of proximal caregiving during threat is touch. Touch is the first of our senses to develop, emerging at around eight weeks gestation^7^. By the third trimester, foetuses differentially respond to their mother’s touch to the abdomen as compared to stranger touch and no touch^8^, setting the stage for one of the earliest maternal interactions. Touch is the primary mode of these interactions, being a necessary part of all caregiving^9,10^. In non-human mammals, it has long being established that touch between conspecifics has evolved to promote not only caregiving but also stress regulation and affiliative bonding^11^, with well-studied neurophysiological, genetic and epigenetic mechanisms^12–14^. Interestingly, idiosyncratic differences in maternal tactile behaviours lead to individual differences in rats’ behavioural and neuroendocrinal responses to stress during adulthood^15^.

There is also increasing understanding in humans about the role of touch in promoting affiliative bonds, affect regulation and healthy development (e.g.^10,16^), while early social and tactile deprivation have corresponding detrimental effects (e.g.,^17,18^). Specifically, following on from the animal literature, research has explored the impact of maternal touch on human infants’ emotion regulation and particularly on stress responses in hypothalamic–pituitary–adrenal (HPA) axis reactivity. Social touch during experimental stress procedures that affect HPA reactivity lead to a decrease in infants’ physiological (e.g., cortisol levels^19^) and behavioural stress responses (e.g., crying), as well as an increase in social attention (eye contact) and positive affect (smiles and vocalizations^20,21^). Moreover, following prenatal depression, the self-reported degree of naturally occurring, maternal stroking touch was capable of mitigating some of the negative effects of the mother’s prenatal depression on infants’ stress reactivity^22^, mirroring similar findings in non-human animals. The effects of touch on cognitive and affective development extend to self-awareness (e.g.,^23,24^) and social learning. For instance, touch is a particularly effective way of directing infant attention^21,25^ and a particularly effective cue for increasing infants’ appropriate eye-contact behaviours^26^.

The effects of touch on brain development were also recently studied in a resting-state functional magnetic resonance imaging study. This investigation revealed a positive relation between the frequency of maternal touch during mother-infant interactions and functional connectivity in various nodes of the infants’ default mode network, thought to support self-awareness and social cognition^16^. Despite this progress in infant research, however, less is known about any lasting effects of such early tactile interactions, and particularly the relationship between individuals’ life-long attachment style and their reactivity to social touch. The primary aim of the present study was, thus, the investigation of this relationship.

We accordingly focused on a neurophysiologically specific type of touch that has been shown to be highly relevant in close relationships, and we investigated how individual differences in adult attachment style may affect its perception. Specifically, while there are many different types of social touch, varying in terms of physiological parameters (e.g., spatial location, temporal properties, pressure), the caress-like, slow velocity, dynamic touch on the skin, known as ‘*affective touch*’ due to its well-studied positive affective value^27^, has been shown to convey social support and intimacy with greater specificity than other types of social touch^28,29^. Critically, this type of touch is thought to convey the rewarding and affiliative properties of touch by means of specialized system of unmyelinated nerve fibres called C-tactile afferents (CTs). CTs are found only in hairy skin^30^, respond only to low pressure stroking and are velocity and temperature tuned; they respond optimally to slow velocity stimulation (1-10cm/s^−1^) and less optimally to velocities below or above this range. The activation of CTs is strongly correlated with perceived pleasantness^31^. Moreover, functional neuroimaging studies on the perception of affective touch have shown selective activation of brain networks that have been associated with the processing of interoceptive signals, that is, signals regarding the physiological condition of the body (i.e., posterior insular cortex^32^, orbitofrontal cortex^33^ and anterior cingulate cortex^34,35^; see also ^36,37^ for reviews and ^38^ for a meta-analysis).

The activation of such networks, dissociations between affective and neutral touch in certain neuropathies^39,40^ and the well-established, positive valence of CT-optimal touch^27,31^ has led some researchers to classify affective touch as a separate tactile interoceptive modality, along with modalities such as pain, itch and temperature perception, which are different from modalities supporting discriminatory (exteroceptive) tactile perception^37,41^. Accordingly, CT-fibers are thought of as the peripheral end of a dedicated interoceptive tactile system supporting the affective and affiliative functions of touch^42,43^. Consistent with these proposals, recent evidence suggests that infants as young as nine months, i.e. during the critical period for the development of IMW, show selective behavioural and physiological sensitivity to stroking touch at ‘CT-optimal’ velocities as compared to similar but ‘non CT-optimal’ velocities (< 1cm/s^−1^ and > 10cm/s^−1^) ^44^. Indeed, the notion that the neural substrate for detecting pleasure associated with this kind of touch develops early has been supported by recent findings suggesting that gentle touch at CT-optimal speeds in newborns leads to increased BOLD activity in brain regions associated with the socio-affective processing of touch (i.e., postcentral gyrus^45^ and insular cortex^45,46^). A recent behavioural study further found that this type of touch helps infants tune to social signals, such as faces, more than other types of touch^47^. These findings support the idea that the CT system may form part of a dedicated interoceptive modality conveying affective and affiliative aspects of social touch and thus promoting the development of self-awareness and affect regulation in early development^24^.

In adults, stroking touch at CT-optimal velocities, as compared with non-optimal velocities, has been shown to specifically communicate social intimacy and support^28^, to reduce experimentally-induced feelings of social rejection^29^ and subjective and neural responses to noxious stimulation^48^, as well as to contribute uniquely to embodied facets of self-awareness^49–51^. Nevertheless, the relationship between CT-optimal stimulation, autonomic regulation, and interoception remains unclear in adults, as CT-optimal touch is specifically associated with reductions in cardiac reactivity and skin conductance responses^52^, but not other measures of autonomic reactivity, such as cortisol variability^53^, or interoceptive awareness such as heart rate awareness^50,53^. Despite the increasing emphasis in the literature on the relationship between interoceptive awareness and emotion regulation, such mixed findings in adult cohorts are not uncommon (see ^54,55^ for discussion). It is indeed increasingly recognized that while sensing of bodily signals (interoception) is a fundamental facet of our emotional experience, the explanatory power of heartbeat detection accuracy only weakly predicts an individual’s vulnerability to anxiety or other affective disorders. Instead, it appears that in order to specify the role of interoception in personality and psychopathology, models need to account for the relationship between bottom-up (e.g., interoceptive accuracy abilities) and top-down (cognitive beliefs, styles, and expectations) factors. Accordingly, this study aimed to characterize individual differences in affective touch and cardiac awareness in terms of differences in pre-existing models of social interactions, namely attachment styles.

Indeed, if affective touch is an interoceptive modality particularly relevant to social affiliation and affect regulation, one can presume that the perception of affective touch in adulthood can further depend on individual differences in attachment. As pre-existing affective-cognitive models of social relating, individual differences in attachment style could determine the top-down influences on the perception of affective touch. Such findings exist in other interoceptive modalities such as hunger^56^ and pain (reviewed by ^57,58^), the latter being an interoceptive modality with opposite hedonic (positive vs. negative valence) and social (care vs. harm) characteristics to affective touch (see^59^ for discussion). For example, chronic pain is more common in individuals with insecure (characterized by early interpersonal experiences of inconsistent and unreliable caregiving) rather than secure attachment styles (characterized by comfort in closeness and consistent, reliable caregiving^57^). We have also previously shown that the effects of social support on subjective, physiological and neural responses to pain, including support conveyed by CT-optimal touch^48^, depend on individual differences in attachment style^48,60,61^.

However, to our knowledge, the relationship between attachment style and sensitivity to affective touch has not yet been studied. Here, we present the first study to examine how attachment style relates to the perception of affective touch, as well as to a different non-social modality of interoception, namely cardiac sensitivity. Specifically, acknowledging the possible continuity of attachment styles from childhood to adulthood and taking into account different research traditions in measuring attachment (see^62^ for a review), we examined attachment using two different measures. First we administered the gold standard Adult Attachment Interview (AAI ^63^). Developed in the context of early, classic research into attachment styles (Main *et al.*, 1985) and thus taking a categorical approach, this semi-structured interview yields secure vs. insecure attachment classifications (as well as further sub-classifications of attachment characteristics such as insecure preoccupied and insecure dismissive) on the basis of questions relating to childhood experiences with caregivers. In addition, we used a well-validated self-report questionnaire (the Experiences in Close Relationships Revised questionnaire^64^). This questionnaire pertains to adult romantic relationships and takes a dimensional rather than categorical approach. In line with theory and research suggesting that adult attachment styles are best conceptualised as dimensional constructs^65^, this questionnaire yields continuous scores of attachment anxiety and avoidance (lower scores denoting greater attachment security and higher scores greater insecurity). Attachment anxiety is characterized by a need for emotional closeness, worries of rejection and abandonment, over-dependence on others, negative views of self, positive views of others, and high emotional reactivity. Attachment avoidance is characterized by a need for emotional distance, resistance to trusting and depending on others, positive views of self, negative views of others, and suppression of emotion.

Given their history in seeking comfort through proximity, we expected that securely attached individuals, based on a categorical AAI classification, would find affective touch (at CT-optimal speeds) more pleasant than non-affective, neutral touch (at non-CT-optimal speeds). By contrast, we expected that insecurely attached individuals (associated with reduced proximity-seeking in the case of dismissive attachment, or truly obtaining comfort through proximity, including touch, in the case of preoccupied attachment) would be less sensitive to affective touch, that is, they would show reduced perceived pleasantness discrimination between the two types of touch. Exploring such differences further using a continuous measure of adult attachment style, we expected that this reduced sensitivity to the hedonic effects of affective (CT-optimal) and neutral (non-CT-optimal) touch would be especially pronounced in individuals scoring higher in anxious and avoidant attachment dimensions, given their typical negative feelings and beliefs about seeking, or receiving social support^66^.

In addition, to investigate whether the relationship between insecure attachment and affective touch sensitivity is specific to this modality, or whether it relates to all interoceptive domains, we also employed a widely-used task of heartbeat counting as a measure of ‘interoceptive accuracy’, a particular facet of interoceptive awareness (see ^67^). Given previous findings about the dissociation between cardiac accuracy and affective touch^50,68^, we expected that attachment style, as measured by both categorical and continuous measures, would not relate to cardiac perceived accuracy as measured by the standard heartbeat perception task, confirming the specificity of affective touch to social bonding and attachment.

## Method

Participants were *N* = 44 right-handed women aged 18 – 31 years old (*M* = 23.87, *SD* = 3.77), recruited from King’s College London and University College London. Participants did not currently suffer from and/or have a history of psychiatric disorders, neurological or medical conditions, and did not have wounds, scars, tattoos or skin irritation/diseases on their forearms. Participants were invited to take part in a study on bodily self-awareness consisting of two separate parts: one part involved rating the pleasantness of touch administered by the experimenter at different velocities (the touch paradigm; see below) and an interoceptive accuracy (heartbeat perception) task. The other part comprised the adult attachment interview, and participants also completed the Experiences in Close Relationships Revised questionnaire (ECR-R^64^) a self-report measure of adult attachment style. Participants’ numerical IDs were used to match data from the different parts of the study and written informed consent was obtained from all participants. The Chair of the Research Department of Clinical, Educational and Health Psychology, University College London (UCL), approved this study and the experiment was conducted in accordance with the Declaration of Helsinki.

### Touch paradigm

A trained experimenter unknown to participants manually stroked participants’ left forearm using a cosmetic make-up brush (Natural hair Blush Brush, No 7, The Boots Company). Participants were seated comfortably at a computer, with their left forearm rested at an approximate 45° angle in front of them (their palm facing upwards) but separated from their view by means of a curtain. Two 9cm long by 4cm wide areas were marked continuously along participants’ left volar forearm between wrist and elbow. To ensure a constant pressure, the brush splayed no wider than the 4cm window. Touch was administered to the underside of participants’ left forearm in an elbow-to-wrist direction^69,70^ at four different velocities, administered in a pseudo-randomized order and alternating between skin areas to avoid habituation: two CT-optimal speeds i.e., 3cms^-1^ and 9cms^−1^ and two non-CT-optimal speeds i.e., 0.3cms^-1^ and 27cms^−1^. Each velocity was administered for 9s, followed by a 30-second interval during which participants rated the pleasantness of the touch on a scale from -100 (very unpleasant) to 100 (very pleasant). Each velocity was administered three times and a mean rating was calculated for each velocity.

### Interoceptive accuracy

We measured interoceptive accuracy using the heartbeat perception task^71^. Participants’ heart rate was recorded using MP150 Data Acquisition Hardware (BIOPAC Systems Inc). A heartbeat monitor was attached to the tip of the left index finger and checked for tightness so that participants could not feel a pulse at this site. During a short training session participants were instructed to report the number of perceived heartbeats within a 15 seconds time interval. They were explicitly told to only count and report the number of actually perceived (and not estimated) heartbeats. The experiment started with a 10 second resting period. Participants closed their eyes and then silently counted their heartbeat (keeping their hand still and without feeling their pulse) for three trials lasting 25 seconds, 35 seconds and 45 seconds; the order was pseudorandomised and participants were not informed of the duration of each trial. The beginning and end of each counting interval was signaled via tones. There was a 20 second pause after each trial during which participants verbally indicated their count for each trial. Interoceptive accuracy was computed using the mean score of the three heartbeat counting trials, using the transformation detailed in ^71^ (see formula also below).

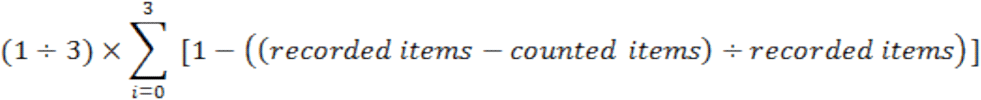

This yields a score between zero and one, with one denoting greater correspondence between actual and perceived number of heartbeats i.e., higher interoceptive accuracy.

### Adult attachment interview (AAI; George et al., 1996)

The AAI is a semi-structured interview, including 20 questions and lasting up to circa one hour. Meta-analyses and psychometric testing indicate stability, and discriminant and predictive validity in both clinical and non-clinical populations^62, 72–74^. Participants were asked to reflect about their childhood experiences and early relationships with parents/caregivers. Questions included whether participants had experienced loss, separation or rejection, how their caregiver typically responded in particular situations e.g., when the participant was upset, and the kinds of implications these experiences had for the participant’s adult life (see^72^ for a detailed introduction to the AAI). All interviews were audio recorded and transcribed verbatim (including pauses). A trained coder (C.V.) coded all interview transcripts and classified participants as secure, dismissive, preoccupied, or unresolved, which allowed us to categorize participants as either securely or insecurely (dismissive, preoccupied, unresolved) attached i.e., our main two categories of interest. A second trained coder (T.N.) independently coded 25% of interviews. Agreement between the two coders was perfect (Cohen’s kappa = 1) for the secure vs. insecure classification. Six participants did not attend the AAI session; hence, *n* = 38 participants were included in these analyses.

### Self-report measure of adult attachment style (Experiences in Close Relationships Revised – ECR-R; Fraley et al., 2000)

The ECR-R comprises 36 items rated on a 7-point scale (1 = strongly disagree and 7 = strongly agree) regarding the general experience of intimate adult relationships; 18 items pertain to attachment anxiety (e.g., “I’m afraid that I will lose my partner’s love.”) and 18 to attachment avoidance (e.g., “I don’t feel comfortable opening up to romantic partners.”). Item responses are averaged (after reverse-scoring appropriate items) separately for each subscale to produce a mean score for attachment anxiety and attachment avoidance, with higher scores denoting greater attachment insecurity. This dimensional scoring is in line with research indicating that adult attachment styles are best conceptualised as dimensional constructs^65^. The ECR-R is well-validated^62,75^ and demonstrates excellent internal consistency: Cronbach’s α = .91 for attachment anxiety and α = .90 for attachment avoidance in the present sample.

### Data availability

The datasets generated during and/or analysed during the current study are available from the corresponding author on reasonable request.

### Results

#### Descriptive statistics and Preliminary Analyses

##### Affective Touch Perception

Pleasantness ratings showed an inverted U-shaped pattern commonly observed for these velocities (see e.g.,^31^ : ratings were lowest for velocities at either end of the velocity spectrum i.e., the 0.3cm/s^−1^ and 27cm/s^−1^ velocities, and highest for the intermediate velocities i.e., 3cm/s^−1^ and 9cm/s^−1^ (see Figure 1). Velocity was associated with pleasantness ratings, *F*(3, 129) = 59.46, *p* < 0.001, with Sidak-corrected pairwise comparisons indicating that velocities differed significantly from each other (*p* < .001) except for the two CT optimal velocities (3cm/s^−1^ vs. 9cm/s^−1^, *p* = .999) and the two non-CT-optimal velocities (0.3cm/s^-1^ vs. 27cm/s^−1^, *p* = .807). Therefore, we computed mean ratings for CT-optimal vs. non-CT-optimal velocities by calculating the average of ratings for 3cms^−1^ and 9cms^−1^ speeds, and 0.3cms^−1^ and 27cms^−1^ speeds, respectively. CT-optimal vs. non-CT-optimal velocities differed as expected, paired samples *t*(43) = 11.41, *p* < .001, with CT-optimal velocities perceived as more pleasant (*M* = 48.43, *SE* = 3.34) than non-CT-optimal velocities (*M* = 3.55, *SE* = 3.74).

**Figure 1.**
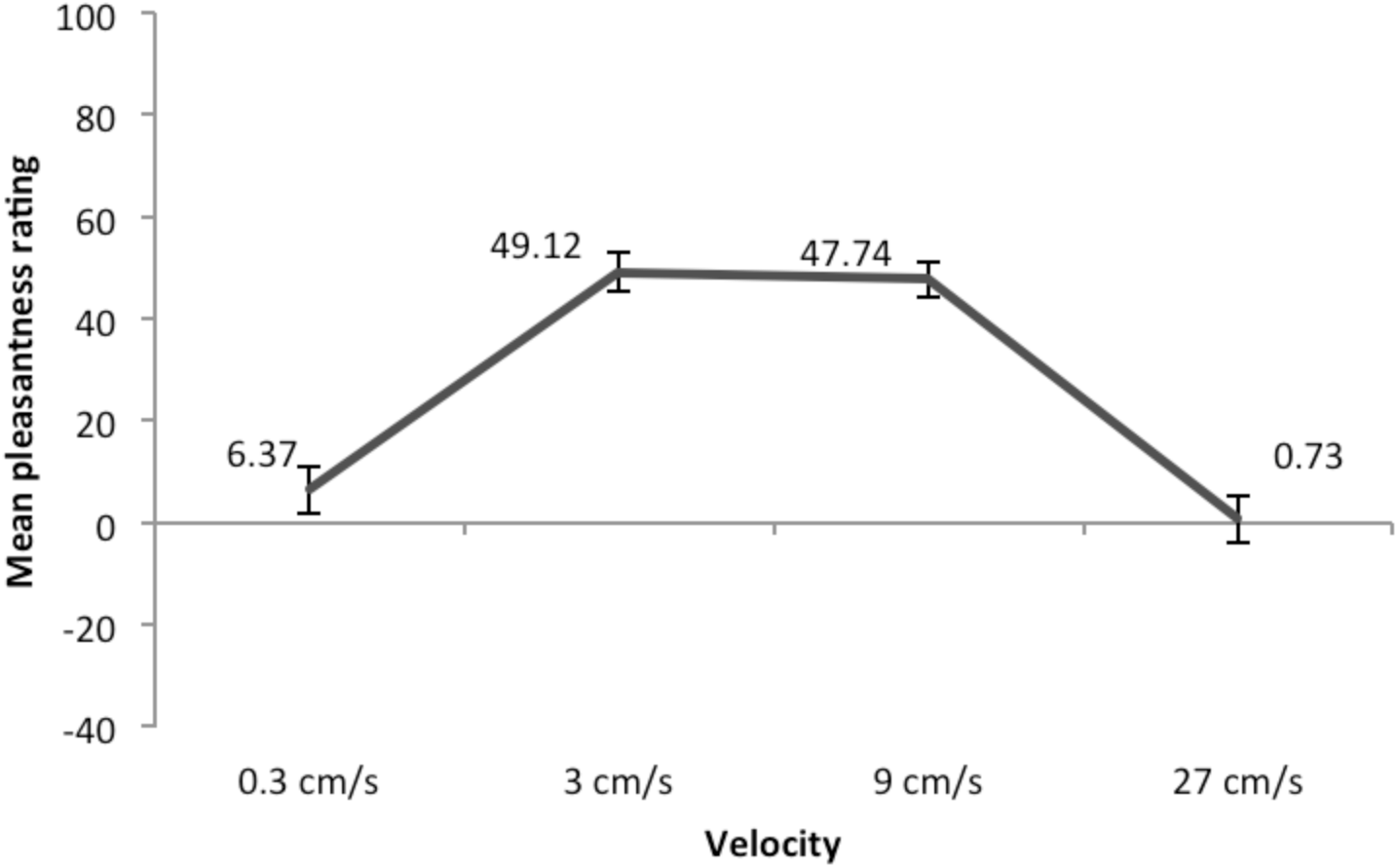
Pleasantness ratings for the four touch velocities. Scale range: −100 to 100. Error bars denote ±SEM.

##### Interoceptive (Cardiac) Accuracy

Mean (*SD*) interoceptive accuracy was .42 (.25). The obtained mean score was slightly lower than mean scores in previous studies using this paradigm (e.g.,^50,67^).

##### Relationship Between Affective-Neutral Touch Discrimination and Interoceptive (Cardiac) Accuracy

Affective-neutral touch discrimination (operationalised as a difference score of CT-optimal velocities minus non-CT-optimal velocities, i.e., greater scores denoting higher pleasantness ratings for CT-optimal vs. non-CT-optimal touch) was not significantly correlated with interoceptive (cardiac) accuracy, *r* = .21, *p* > .05.

##### Attachment Classifications and Dimensions

Based on the AAI, our sample (*n* = 38) showed the following classification frequencies: *n* = 30 participants (79%) were classified as securely attached and *n* = 8 (21%) as insecurely attached, of which *n* = 1 (3%) was preoccupied, *n* = 5 (13%) were dismissing, and *n* = 2 (5%) were unresolved. Despite an overrepresentation of securely attached individuals, our sample is in line with the general population AAI norms for non-clinical adult mothers^73^ as well as non-clinical adolescents/students^76^, also suggesting a larger proportion of dismissive vs. preoccupied individuals. Mean (*SD*) ECR-R dimensional anxiety scores = 2.99 (0.98) and avoidance scores = 3.03 (0.93); in relation to general population norms for women, our sample fell below the mean for anxiety (population norm *M* = 3.56, *SD* = 1.13) and above the mean for avoidance (population norm *M* = 2.92, *SD* = 1.21; see information by Fraley, 2012: http://internal.psychology.illinois.edu/~rcfraley/measures/ecrr.htm). ECR-R dimensions were moderately correlated with each other, *r* = .49, *p* < .001, and were mean centered in statistical analyses to minimize multicollinearity issues^77^.

##### The Relationship between Categorical and Dimensional Measures of Attachment Style

A MANOVA with ECR-R anxiety and ECR-R avoidance scores as outcome variables and AAI classification (secure vs. insecure) as the independent variable showed that ECR-R scores did not differ by AAI classification, *F*(2, 36) = 1.74, *p* = .191, Wilk’s lamda = .910. In other words, it was not the case that ECR-R anxiety scores were significantly lower in the secure vs. insecure group (Secure: *M* = 2.87, *SD* = 0.98; Insecure: *M* = 3.37, *SD* = 0.78), or that ECR-R avoidance scores were significantly lower for securely vs. insecurely attached participants (Secure: *M* = 3.05, *SD* = 1.00; Insecure: *M* = 2.90, *SD* = 0.71). This result supports the choice of two separate measures for this multi-dimensional construct.

##### The Relationship between Measures of Attachment and Interoceptive Accuracy

AAI classification was not associated with interoceptive accuracy, *F*(1,35) = 0.45, *p* = .505. In addition, neither ECR-R anxiety (*r* = −.05, *p* > .05) nor ECR-R avoidance scores (*r* = .13, *p* > .05), were significantly correlated with interoceptive accuracy, as predicted.

### Main Analyses

#### Association between interview-assessed attachment style (AAI classification) and the perception of affective touch

To examine whether attachment classification as measured by the AAI was associated with the perception of affective touch, we specified a multilevel regression model with mean pleasantness rating as the outcome variable and velocity (CT-optimal vs. non-CT-optimal), AAI (security vs. insecurity), and their interaction as predictor variables, and controlled for interoceptive accuracy. A random effect was included to account for the repeated assessment of the outcome variable within individuals.

AAI classification predicted pleasantness ratings across velocities: insecurely attached participants rated touch as more pleasant (*M* = 34.52, *SE* = 7.46) than did securely attached participants (*M* = 23.21, *SE* = 3.59). More critically, the hypothesised velocity by AAI interaction was significant (see Table 1 for full model results). Follow-up tests showed that the difference between CT-optimal and non-CT-optimal velocities was significant for securely attached participants (*b* = 49.01, *SE* = 4.08, *p* < .001), and insecurely attached participants (*b* = 28.19, *SE* = 8.44, *p* = .001). However, the difference in pleasantness ratings for CT-optimal and non-CT-optimal velocities was smaller for insecurely vs. securely attached participants (see adjusted mean difference above and Figure 2, top panel): an independent samples t-test on the affective-neutral touch difference score (see above for how this was computed) confirmed that the difference was smaller in the insecure group (*M* = 28.81, *SD* = 14.13) than the secure group (*M* = 49.01, *SD* = 4.85), *t*(36) = 2.06, *p* = .047. Therefore, although both groups were able to discriminate between the two forms of touch, attachment insecurity was associated with reduced discrimination between CT-optimal and non-CT-optimal touch, in line with our hypothesis.

**Table 1.** Model results for effects of attachment on the perception of affective touch.

|  |  | *b* | *SE* | *p* value | 95% CI<br>lower | 95% CI<br>higher |
| --- | --- | --- | --- | --- | --- | --- |
| AAI | Velocity (CT-optimal vs. Non-CT-optimal) | 49.01 | 4.08 | < **0.001** | 41.12 | 57.00 |
|  | AAI (security vs. insecurity) | 21.72 | 9.53 | **0.023** | 3.05 | 40.40 |
|  | Velocity by AAI | -20.82 | 9.37 | **0.026** | -39.19 | -2.45 |
| ECR-R | Velocity (CT-optimal vs. Non-CT-optimal) | 44.30 | 3.53 | < **0.001** | 37.39 | 51.21 |
|  | ECR-R anxiety | 5.04 | 3.95 | 0.202 | -2.71 | 12.78 |
|  | ECR-R avoidance | -3.92 | 4.21 | 0.352 | -12.16 | 4.33 |
|  | Velocity by ECR-R anxiety | -8.93 | 3.90 | **0.022** | -16.58 | -1.28 |
|  | Velocity by ECR-R avoidance | -2.09 | 4.14 | 0.614 | -10.20 | 6.02 |
|  | ECR-R anxiety by ECR-R avoidance | -0.25 | 2.88 | 0.932 | -5.89 | 5.40 |
|  | Velocity by ECR-R anxiety by ECR-R avoidance | 1.27 | 2.86 | 0.656 | -4.33 | 6.88 |

*Note.* AAI = Adult Attachment Interview; ECR-R = Experiences in Close Relationships – Revised questionnaire.

**Figure 2.**
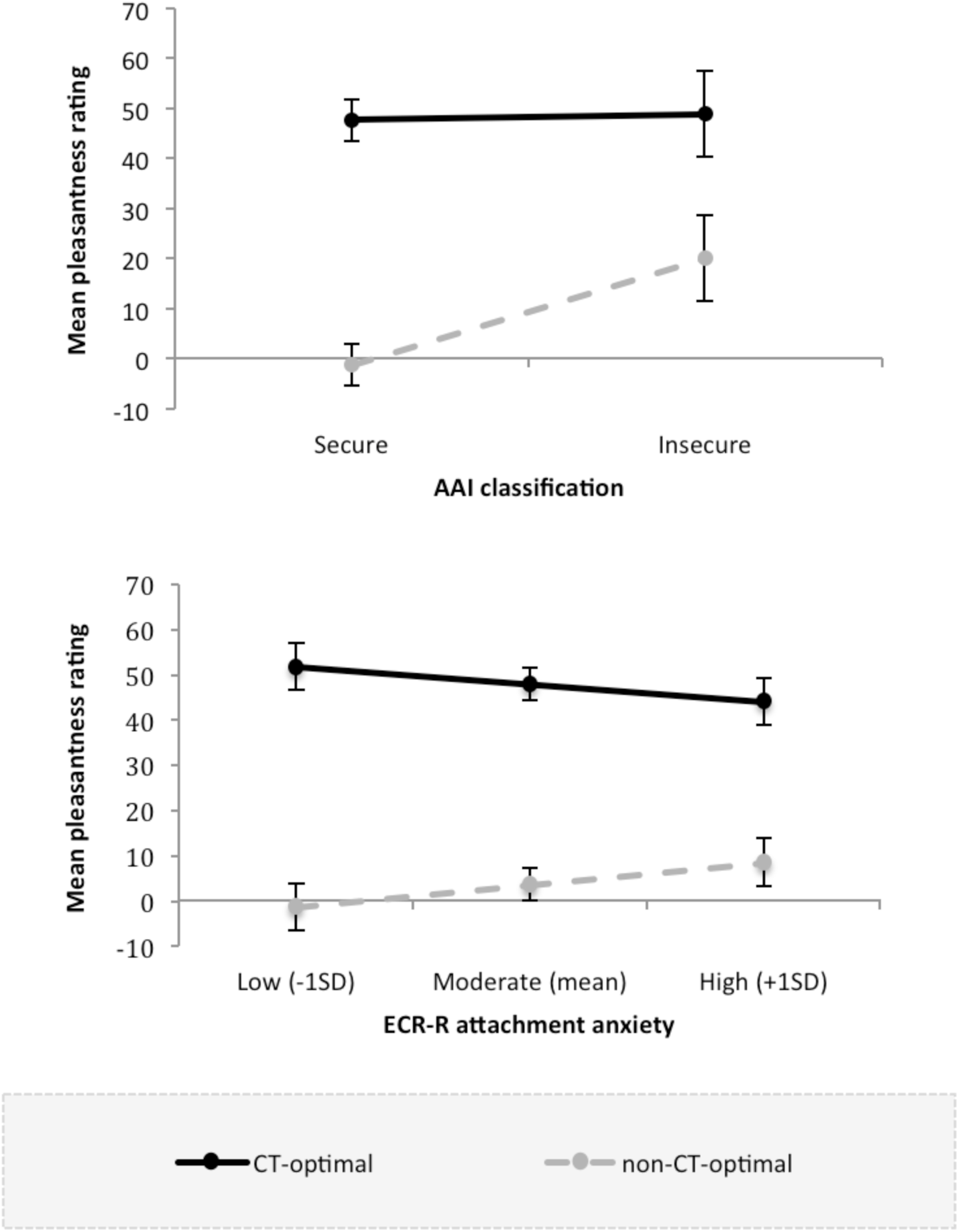
Interactions between velocity (CT-optimal vs. non-CT-optimal) and attachment classification on the Adult Attachment Interview (AAI; top panel), and velocity and attachment anxiety measured using the Experiences in Close Relationships – Revised questionnaire (ECR-R; bottom panel). Error bars represent ±1 standard error of the mean.

#### Association between questionnaire-assessed attachment style (ECR-R) and the perception of affective touch

To test whether attachment style dimensions as measured by the ECR-R questionnaire were associated with the perception of pleasant touch, we specified a multilevel regression model with mean pleasantness rating as the outcome variable, and velocity (CT-optimal vs. non-CT-optimal), ECR-R attachment anxiety, ECR-R attachment avoidance, as well as all interaction terms, as predictor variables, and again controlled for interoceptive accuracy. As above, a random effect was included to account for the repeated assessment of the outcome variable within individuals.

Neither attachment anxiety nor attachment avoidance, nor the interaction between the two dimensions were associated with pleasantness ratings across velocities, indicating that pleasantness of touch in general was not influenced by continuous attachment style scores. However, importantly, the hypothesised velocity by attachment anxiety interaction was significant (see Table 1 for full model results). Follow-up analyses revealed that the difference between CT-optimal and non-CT-optimal velocities was significant at lower (i.e., −1SD; *b* = − 53.06, *SE* = 5.12, *p* < .001), moderate (i.e., mean; *b* = −44.32, *SE* = 3.52, *p* < .001) and higher (i.e., +1SD; *b* = −35.58, *SE* = 5.24, *p* < .001) levels of attachment anxiety. Similar to the AAI results, the difference in pleasantness ratings between CT-optimal and non-CT-optimal velocities was smallest at higher levels of attachment anxiety (see adjusted mean difference above and Figure 2, bottom panel). This finding indicates that higher attachment anxiety was associated with reduced discrimination between CT-optimal and non-CT-optimal touch. The velocity by attachment avoidance interaction was non-significant, as was the three-way velocity by attachment anxiety by attachment avoidance interaction. Thus, attachment avoidance was not associated with the perception of affective touch, either alone or interaction with attachment anxiety. In sum, partially supporting our hypothesis, higher attachment anxiety but not attachment avoidance was associated with reduced sensitivity to affective touch.

## Discussion

We reported the first study investigating the association between attachment styles and affective touch in adulthood. Under the assumption that affective touch supports affiliative bonds and social cognition, we assessed how affective-cognitive models of social relating (i.e., attachment) influence the perception of affective touch. Using the gold standard assessment of adult attachment (the Adult Attachment Interview; AAI), we found that insecure attachment was associated with reduced discrimination between affective vs. non-affective, neutral touch. This semi-structured interview yields categorical attachment classifications in an implicit way, relating to representations of childhood experiences with caregivers. Acknowledging the different traditions in measuring attachment and the multi-dimensionality of this construct, we also used a well-validated self-report questionnaire that pertains to explicit evaluations of close relationships and takes a dimensional rather than categorical approach. This measure showed that higher attachment anxiety, though not higher attachment avoidance, was associated with reduced discrimination between affective vs. non-affective, neutral touch. Attachment style as assessed by both measures was not related to cardiac perception accuracy, suggesting that attachment is not relevant to all interoceptive modalities in the same way. These results will be discussed in turn below.

We found that both secure and insecure attachment groups (assessed by the AAI) were able to discriminate between affective vs. non-affective touch. However, the insecure attachment group was significantly worse in this discrimination than the secure group, suggesting that differences in pre-existing models of social interaction and related top-down expectations contribute to individual differences in affective touch perception. This finding provides further support to theoretical proposals regarding an association between the perceived affectivity of touch and social affiliation^24,42,43,59^. Attachment representations are thought to originate in early caregiving experiences, in which touch plays a central part^9,10^. There is also evidence that childhood patterns of social relationships may be reinforced across the lifespan^5^ and it thus appears that affective responses to touch are also carried into adulthood.

Differences in attachment (as measured by the AAI) were not related to cardiac accuracy. To date, there is no evidence to suggest that cardiac awareness is an interoceptive modality relevant to social affiliation and thus pre-existing models of social relating, such as attachment classifications, were not predicted to influence this interoceptive modality. This finding also speaks to the more general relationship between interoceptive modalities. In other studies, our group and others have shown that the perception of social, affective touch and cardiac perception accuracy are unrelated^50,68^. The current findings confirm this and further suggest that differences in attachment relate to the perception of affective touch but not cardiac perception accuracy.

Turning to our second two-dimensional measure of attachment style, we found that higher scores in attachment anxiety were related to poorer discrimination between affective and non-affective touch. This finding suggests that anxious attachment style as assessed by an explicit measure of adult close relationships relates to the perceived affectivity of touch in a similar way to insecure attachment as assessed by the AAI. In insecure attachment, others are perceived as unreliable and inattentive, and particularly in anxious insecure attachment this might generate anxiety^78^. These kinds of social expectations might thus affect the way in which affective touch is perceived and enjoyed. Consistent with recent research suggesting that infants as old as two months old show selective sensitivity to affective touch^45,46^, we hypothesized that any difference in this discrimination in adulthood based on attachment styles would relate to top-down effects. Indeed, we also observed that insecure attachment style was related to the overall perceived pleasantness of our tactile stimuli, irrespective of whether or not they were in the CT-optimal range.

The fact that we did not observe differences in discrimination in our questionnaire dimension of attachment avoidance was unexpected, particularly as these individuals are characterized by a need for emotional distance and reduced proximity-seeking, including touch^62,79,80^. This null finding thus suggests that at least at an explicit level, current top-down representations of close relationships in these individuals may not determine the perceived affectivity of touch. However, future research is needed before drawing firm conclusions. Here, we propose a few candidate explanations that may have led to such a lack of findings. First, although measurements of attachment style may possess benefits at a theoretical and statistical level, self-reported questionnaires have been largely criticized for being passive, i.e., not detecting attachment phenomena that need to be activated to be manifested ^62^. As such, this may have contributed to the current lack of findings. Also, as self-report measures, they are subject to social desirability effects, which are likely to be more pronounced in more avoidantly-attached individuals.

Finally, as with our other implicit measure of attachment, we found that individual differences in attachment style (as measured by the questionnaire dimensions) were not related to cardiac accuracy, suggesting that cognitive more models of current close social relationships and related top-down expectations do not contribute to individual differences in cardiac interoceptive perception. Given that individual differences in attachment style were related to the perceived affectivity of the touch and not cardiac accuracy, this finding provides further support to the specificity of the relationship between affective touch and attachment style.

Our findings should be considered in light of their limitations and directions for future research. First, it should be noted that there were no differences in the attachment anxiety or avoidance questionnaire scores between the secure and insecure AAI groups. This finding, together with prior research suggesting a trivial to small relation between self-report measures of attachment and the AAI^81^, speaks to the different aspects captured by each of these measures and consequently supports the choice of two separate measures for this multi-dimensional construct. Second, on the AAI, small numbers in the insecure attachment group meant we were unable to further compare preoccupied vs. dismissing individuals. Future studies could aim to recruit larger groups of preoccupied and dismissing individuals to examine whether results on the insecure group may have been driven by preoccupied or dismissive individuals (although interestingly, the largest subgroup in the insecure AAI classification was dismissive; in line with the general population AAI norms^73,76^). Third, pleasantness ratings for the affective touch velocities overall fell in the middle of the positive side of the response scale; it is likely that touch by an attachment figure, such as the romantic partner, may feel even more pleasant to participants than touch by an experimenter. Similarly, the current effects on affective vs. non-affective touch discrimination may be subject to social context, in which for instance, touch by an attachment figure could activate attachment behaviors that are not always at display (see ^62^). Thus, future research could incorporate partner-administered touch. Finally, we only tested women in order to control for gender effects associated with the perception of touch ^70,82,83^; however, future research is needed to examine whether the present results extend to men.

In sum, the present study corroborates and extends previous literature on the affectivity of touch and its relation with affiliative bonds and social cognition. Given that attachment style (in both measures) was not related to perceived cardiac accuracy, these findings point to the specificity of the relationship between affective touch and attachment style. Future work is needed to examine the role of social context and whether the present results extend to men.

## Acknowledgements

We are grateful to Nadia Blom, Amanda Hornsby and Isabel Rafferty for their help in transcribing the adult attachment interviews, and to Nikolaos Tzikas and Elena Panagiotopoulou for his help in conducting several of the interviews. This study was supported by a project grant (no. II/85 069) from the Volkswagen Foundation ‘European Platform for Life Sciences, Mind Sciences and Humanities’ and a European Research Council Starting Investigator Award (ERC-2012-STG GA313755) (to A.F.). A.F.’s time was also partly supported by an ApsaA research grant. T.N. is supported by the Wellcome Trust Principal Research Fellowship of Professor Read Montague. M.V. M. is supported by the National Council on Science and Technology (CONACyT- 538843; Mexico) scholarship.

## Author Notes

### Author contributions

C. Krahé, A. Gentsch and A. Fotopoulou developed the hypothesis and research plan. L. Guy collected the data. C. Vari and T. Nolte coded the interviews. C. Krahé and M. Von Mohr analyzed the data. C. Krahé and M. Von Mohr wrote the manuscript, under the guidance of A. Gentsch, C. Vari, T. Nolte, and A. Fotopoulou. All authors approved the final version of the manuscript for submission.

## References

1. Bowlby, J. The making and breaking of affectional bonds: Aetiology and psychopathology in light of attachment theory. Br. J. Psychiatry 130, 201–10 (1977).

2. Bowlby, J. Attachment and loss: Attachment. Attachment 1, (1969).

3. Main, M., Kaplan, N. & Cassidy, J. Security in Infancy, Childhood, and Adulthood: A Move to the Level of Representation. Monogr. Soc. Res. Child Dev. 50, 66 (1985).

4. Hazan, C. & Shaver, P. Romantic love conceptualized as an attachment process. J. Pers. Soc. Psychol. 52, 511–524 (1987).

5. Waters, E., Merrick, S., Treboux, D., Crowell, J. & Albersheim, L. Attachment security in infancy and early adulthood: a twenty-year longitudinal study. Child Dev. 71, 684–9 (2000).

6. Mikulincer, M., Shaver, P. R., Sapir-Lavid, Y. & Avihou-Kanza, N. What’s inside the minds of securely and insecurely attached people? The secure-base script and its associations with attachment-style dimensions. J. Pers. Soc. Psychol. 97(4), (2009).

7. Montagu, A. Touching: The human significance of the skin. (Harper & Row, 1978).

8. Marx, V. & Nagy, E. Fetal behavioral responses to the touch of the mother’s abdomenA Frame-by-frame analysis. Infant Behav. Dev. 47, 83–91 (2017).

9. Stack, D. M. in Blackwell Handbook of Infant Development 351–378 (2007). doi:10.1002/9780470996348.ch13

10. Field, T. Touch for socioemotional and physical well-being: A review. Developmental Review 30, 367–383 (2010).

11. Dunbar, R. I. M. The social role of touch in humans and primates: Behavioural function and neurobiological mechanisms. Neuroscience and Biobehavioral Reviews 34, 260–268 (2010).

12. Harlow, H. F. & Harlow, M. Social deprivation in monkeys. Sci. Am. 207, 136–146 (1962).

13. Nelson, E. E. & Panksepp, J. Brain substrates of mother-infant attachment: Contributions of opoids, oxytocin and norepinephrine. Neurosci. Biobehav. Rev. 22, 437–452 (1998).

14. Weaver, I. C. G. et al. Epigenetic programming by maternal behavior. Nat. Neurosci. 7, 847–854 (2004).

15. Zhang, T. Y., Chretien, P., Meaney, M. J. & Gratton, A. Influence of naturally occurring variations in maternal care on prepulse inhibition of acoustic startle and the medial prefrontal cortical dopamine response to stress in adult rats. J Neurosci 25, 1493–1502 (2005).

16. Brauer, J., Xiao, Y., Poulain, T., Friederici, A. D. & Schirmer, A. Frequency of Maternal Touch Predicts Resting Activity and Connectivity of the Developing Social Brain. Cereb. Cortex 26, 3544–3552 (2016).

17. Carlson, M. & Earls, F. Psychological and neuroendocrinological sequelae of early social deprivation in institutionalized children in Romania. in Annals of the New York Academy of Sciences 807, 419–428 (1997).

18. Beckett, C. et al. Do the effects of early severe deprivation on cognition persist into early adolescence? Findings from the English and Romanian adoptees study. Child Dev. 77, 696–711 (2006).

19. Feldman, R., Singer, M. & Zagoory, O. Touch attenuates infants’ physiological reactivity to stress. Dev. Sci. 13, 271–278 (2010).

20. Stack, D. M. & Muir, D. W. Tactile stimulation as a component of social interchange: New interpretations for the still-face effect. Br. J. Dev. Psychol. 8, 131–145 (1990).

21. Stack, D. M. & Muir, D. W. Adult Tactile Stimulation during Face-to-Face Interactions Modulates Five-Month-Olds’ Affect and Attention. Child Dev. 63, 1509–1525 (1992).

22. Sharp, H. et al. Frequency of Infant Stroking Reported by Mothers Moderates the Effect of Prenatal Depression on Infant Behavioural and Physiological Outcomes. PLoS One 7, (2012).

23. Filippetti, M. L., Orioli, G., Johnson, M. H. & Farroni, T. Newborn Body Perception: Sensitivity to Spatial Congruency. Infancy 20, 455–465 (2015).

24. Fotopoulou, A. & Tsakiris, M. Mentalizing homeostasis: The social origins of interoceptive inference-replies to Commentaries. Neuropsychoanalysis 19, 71–76 (2017).

25. Cascio, C. J. Somatosensory processing in neurodevelopmental disorders. J. Neurodev. Disord. 2, 62–69 (2010).

26. Peláez-Nogueras, M. et al. Infants’ preference for touch stimulation in face-to-face interactions. J. Appl. Dev. Psychol. 17, 199–213 (1996).

27. Walker, S. C., Trotter, P. D., Swaney, W. T., Marshall, A. & Mcglone, F. P. C-tactile afferents: Cutaneous mediators of oxytocin release during affiliative tactile interactions? Neuropeptides 64, 27–38 (2017).

28. Kirsch, L. P. et al. Reading the mind in the touch: Neurophysiological specificity in the communication of emotions by touch. Neuropsychologia (2017). doi:10.1016/j.neuropsychologia.2017.05.024

29. Von Mohr, M., Kirsch, L. P. & Fotopoulou, A. The soothing function of touch: Affective touch reduces feelings of social exclusion. Sci. Rep. 7, (2017).

30. Johansson, R. S. & Vallbo, A. B. Tactile sensibility in the human hand: relative and absolute densities of four types of mechanoreceptive units in glabrous skin. J. Physiol. 286, 283–300 (1979).

31. Löken, L. S., Wessberg, J., Morrison, I., McGlone, F. & Olausson, H. Coding of pleasant touch by unmyelinated afferents in humans. Nat. Neurosci. 12, 547–548 (2009).

32. Bjornsdotter, M., Loken, L., Olausson, H., Vallbo, A. & Wessberg, J. Somatotopic Organization of Gentle Touch Processing in the Posterior Insular Cortex. J. Neurosci. 29, 9314–9320 (2009).

33. Mcglone, F. et al. Touching and feeling: Differences in pleasant touch processing between glabrous and hairy skin in humans. Eur. J. Neurosci. 35, 1782–1788 (2012).

34. Lindgren, L. et al. Pleasant human touch is represented in pregenual anterior cingulate cortex. Neuroimage 59, 3427–3432 (2012).

35. Case, L. K. et al. Encoding of Touch Intensity But Not Pleasantness in Human Primary Somatosensory Cortex. J. Neurosci. 36, 5850–5860 (2016).

36. Craig, A. D. How do you feel - now? The anterior insula and human awareness. Nature Reviews Neuroscience 10, 59–70 (2009).

37. Björnsdotter, M., Morrison, I. & Olausson, H. Feeling good: On the role of C fiber mediated touch in interoception. Experimental Brain Research 207, 149–155 (2010).

38. Morrison, I. ALE meta-analysis reveals dissociable networks for affective and discriminative aspects of touch. Hum. Brain Mapp. 37, 1308–1320 (2016).

39. Olausson, H. et al. Unmyelinated tactile afferents signal touch and project to insular cortex. Nat. Neurosci. 5, 900–904 (2002).

40. Olausson, H. et al. Functional role of unmyelinated tactile afferents in human hairy skin: Sympathetic response and perceptual localization. Exp. Brain Res. 184, 135–140 (2008).

41. Craig, a. D. & Craig, a. D. How do you feel? Interoception: the sense of the physiological condition of the body. Nat. Rev. Neurosci. 3, 655–666 (2002).

42. McGlone, F., Wessberg, J. & Olausson, H. Discriminative and Affective Touch: Sensing and Feeling. Neuron 82, 737–755 (2014).

43. Gentsch, A., Crucianelli, L., Jenkinson, P. & Fotopoulou, A. in Affective Touch and the Neurophysiology of CT Afferents 355–384 (2016). doi:10.1007/978-1-4939-6418-5_21

44. Fairhurst, M. T., Löken, L. & Grossmann, T. Physiological and Behavioral Responses Reveal 9-Month-Old Infants’ Sensitivity to Pleasant Touch. Psychol. Sci. 25, 1124–1131 (2014).

45. Tuulari, J. J. et al. Neural correlates of gentle skin stroking in early infancy. Developmental Cognitive Neuroscience (2017). doi:10.1016/j.dcn.2017.10.004

46. Jönsson, E. H. et al. Affective and non-affective touch evoke differential brain responses in 2-month-old infants. Neuroimage 169, 162–171 (2018).

47. Della Longa, L., Gliga, T. & Farroni, T. Tune to touch: Affective touch enhances learning of face identity in 4-month-old infants. Developmental Cognitive Neuroscience (2017). doi:10.1016/j.dcn.2017.11.002

48. Krahé, C., Drabek, M. M., Paloyelis, Y. & Fotopoulou, A. Affective touch and attachment style modulate pain: a laser-evoked potentials study. Philos. Trans. R. Soc. B Biol. Sci. 371, 20160009 (2016).

49. Panagiotopoulou, E., Filippetti, M. L., Tsakiris, M. & Fotopoulou, A. Affective Touch Enhances Self-Face Recognition during Multisensory Integration. Sci. Rep. 7, (2017).

50. Crucianelli, L., Krahé, C., Jenkinson, P. M. & Fotopoulou, A. Interoceptive ingredients of body ownership. Cortex (2017).

51. Crucianelli, L., Metcalf, N. K., Fotopoulou, A. & Jenkinson, P. M. Bodily pleasure matters: Velocity of touch modulates body ownership during the rubber hand illusion. Front. Psychol. 4, (2013).

52. Pawling, R., Cannon, P. R., McGlone, F. P. & Walker, S. C. C-tactile afferent stimulating touch carries a positive affective value. PLoS One 12, (2017).

53. Triscoli, C., Olausson, H., Sailer, U., Ignell, H. & Croy, I. CT-optimized skin stroking delivered by hand or robot is comparable. Front. Behav. Neurosci. 7, (2013).

54. Critchley, H. D. & Garfinkel, S. N. Interoception and emotion. Current Opinion in Psychology 17, 7–14 (2017).

55. Garfinkel, S. N. et al. Interoceptive dimensions across cardiac and respiratory axes. Philos. Trans. R. Soc. B Biol. Sci. 371, 20160014 (2016).

56. Alexander, K. E. & Siegel, H. I. Perceived hunger mediates the relationship between attachment anxiety and emotional eating. Eat. Behav. 14, 374–377 (2013).

57. Meredith, P., Ownsworth, T. & Strong, J. A review of the evidence linking adult attachment theory and chronic pain: Presenting a conceptual model. Clinical Psychology Review 28, 407–429 (2008).

58. Meredith, P. J. A review of the evidence regarding associations between attachment theory and experimentally induced pain. Current pain and headache reports 17, 326 (2013).

59. von Mohr, M. & Fotopoulou, A. in The interoceptive mind: from homeostasis to awareness (eds. Tsakiris, M. & De Preester, H.) (2018).

60. Sambo, C. F., Howard, M., Kopelman, M., Williams, S. & Fotopoulou, A. Knowing you care: Effects of perceived empathy and attachment style on pain perception. Pain 151, 687–693 (2010).

61. Hurter, S., Paloyelis, Y., Amanda, A. C. & Fotopoulou, A. Partners’ empathy increases pain ratings: Effects of perceived empathy and attachment style on pain report and display. Journal of Pain 15, 934–944 (2014).

62. Ravitz, P., Maunder, R., Hunter, J., Sthankiya, B. & Lancee, W. Adult attachment measures: A 25-year review. Journal of Psychosomatic Research 69, 419–432 (2010).

63. George, C., Kaplan, N. & Main, M. Adult attachment interview protocol. Unpubl. manuscript, Univ. Calif. Berkeley (1996).

64. Fraley, R. C., Waller, N. G. & Brennan, K. A. An item response theory analysis of self-report measures of adult attachment. J. Pers. Soc. Psychol. 78, 350–365 (2000).

65. Chris Fraley, R., Hudson, N. W., Heffernan, M. E. & Segal, N. Are adult attachment styles categorical or dimensional? A taxometric analysis of general and relationship-specific attachment orientations. J. Pers. Soc. Psychol. 109, 354–368 (2015).

66. Mikulincer, M., Shaver, P. R. & Pereg, D. Attachment Theory and Affect Regulation : The Dynamics, Development, and Cognitive Consequences of Attachment-Related Strategies 1. Motiv. Emot. 27, 77–102 (2003).

67. Garfinkel, S. N., Seth, A. K., Barrett, A. B., Suzuki, K. & Critchley, H. D. Knowing your own heart: Distinguishing interoceptive accuracy from interoceptive awareness. Biol. Psychol. 104, 65–74 (2015).

68. Triscoli, C., Croy, I., Steudte-Schmiedgen, S., Olausson, H. & Sailer, U. Heart rate variability is enhanced by long-lasting pleasant touch at CT-optimized velocity. Biol. Psychol. 128, 71–81 (2017).

69. Löken, L. S., Evert, M. & Wessberg, J. Pleasantness of touch in human glabrous and hairy skin: Order effects on affective ratings. Brain Res. 1417, 9–15 (2011).

70. Essick, G. K. et al. Quantitative assessment of pleasant touch. Neuroscience and Biobehavioral Reviews 34, 192–203 (2010).

71. Schandry, R. Heart Beat Perception and Emotional Experience. Psychophysiology 18, 483–488 (1981).

72. Hesse, E. The adult attachment interview: protocol, method of analysis, and empirical studies. Handb. Attach. theory, Res. Clin. Appl. 552–598 (2008).

73. van Ijzendoorn, M. H. & Bakermans-Kranenburg, M. J. in Clinical applications of the adult attachment interview 69–96 (2008).

74. Bakermans-Kranenburg, M. J. & Van IJzendoorn, M. H. A psychometric study of the Adult Attachment Interview: Reliability and discriminant validity. Dev. Psychol. 29, 870–879 (1993).

75. Sibley, C. G., Fischer, R. & Liu, J. H. Reliability and validity of the revised Experiences in Close Relationships (ECR-R) self-report measure of adult romantic attachment. Personal. Soc. Psychol. Bull. 31, 1524–1536 (2005).

76. Bakermans-Kranenburg, M. & van IJzendoorn, M. H. The first 10,000 Adult Attachment Interviews: Distributions of adult attachment representations in clinical and non-clinical groups. Attach. Hum. Dev. 11, 223–263 (2009).

77. Aiken, L. S. & West, S. G. Multiple regression: Testin and interpreting interactions. Multiple regression: Testing and interpreting interactions (1991).

78. Nolte, T., Guiney, J., Fonagy, P., Mayes, L. C. & Luyten, P. Interpersonal Stress Regulation and the Development of Anxiety Disorders: An Attachment-Based Developmental Framework. Front. Behav. Neurosci. 5, (2011).

79. Bartholomew, K. & Horowitz, L. M. Attachment styles among young adults: A test of a four-category model. J. Pers. Soc. Psychol. 61, 226–244 (1991).

80. Brennan, K., Clark, C. & Shaver, P. Self-report measurement of adult attachment. Attach. theory close … 46–76 (1998).

81. Roisman, G. I. et al. The Adult Attachment Interview and Self-Reports of Attachment Style: An Empirical Rapprochement. J. Pers. Soc. Psychol. 92, 678–697 (2007).

82. Suvilehto, J. T., Glerean, E., Dunbar, R. I. M., Hari, R. & Nummenmaa, L. Topography of social touching depends on emotional bonds between humans. Proc. Natl. Acad. Sci. 112, 13811–13816 (2015).

83. Gazzola, V. et al. Primary somatosensory cortex discriminates affective significance in social touch. Proc. Natl. Acad. Sci. 109, E1657–E1666 (2012).

